# Unraveling Tyrosine-Kinase Inhibitor Resistance in NSCLC Cells via Same-cell Measurement of RNA, Protein, and Morphological Responses

**DOI:** 10.1101/2025.05.06.652479

**Authors:** Vivian Dien, Connor Thompson, Tristin Rammel, Jaime Roquero, Marielle Krivit, Grace Yeo, Daniel Honigfort, Pazinah Bhadha, Colin Brown-Greeves, Mariam Dawood, Keerthana Elango, Hyukjin J. Kwon, Tyler Lopez, Alvin Lo, Meghna Manjunath, Pin Ren, Ben Song, Mitch Sundkamp, Ramreddy Tippana, Wilfred Van IJcken, Jennifer Wong, Jane Yao, Da Zhou, Sinan Arslan, Matthew Kellinger, Amirali Kia, Semyon Kruglyak, Michael Previte, Molly He

## Abstract

Non-small cell lung cancer (NSCLC) frequently develops resistance to tyrosine kinase inhibitors (TKIs), limiting the long-term success of targeted therapies. A deeper understanding of resistance mechanisms at the molecular and cellular levels may enable the development of more effective treatment strategies. Here, we applied the Teton detection assay on the AVITI24 platform to measure concurrently RNA, protein, and cellular morphology in NSCLC cell lines treated with the TKIs gefitinib and osimertinib. This single-cell, multiomic analysis revealed distinct expression and morphological profiles between drug-sensitive and resistant cells, including differences in MAPK-related pathway activity. Stratifying responses at the single-cell level uncovered subtle responses not detectable in bulk measurements. We identified CDK4/6 activity as a route of cell survival under TKI treatment and demonstrated that co-treatment with the CDK4/6 inhibitor palbociclib enhanced TKI efficacy. The ability to measure multiomics and cellular morphology in the same cells opens new avenues for future studies aimed at improving personalized treatment strategies in NSCLC and overcoming the obstacles posed by drug resistance.

## Introduction

Targeted therapies have transformed the treatment landscape for non-small cell lung cancer (NSCLC), yet the development of resistance remains a significant obstacle to long-term patient benefit. Cell culture systems have been instrumental in understanding drug response mechanisms and advancing therapeutic strategies, from the pioneering in vitro models of the early 20th century^1,2^ to today’s more sophisticated platforms. While conventional cell-based assays provide valuable quantitative insights into drug effects^3–5^, they often fail to capture the complexity of in vivo responses, leading to discrepancies between preclinical promise and clinical reality^6–8^. With fewer than 10% of drug candidates advancing through clinical trials, and many failures due to lack of efficacy or unforeseen toxicity, there is a pressing need for more predictive, biologically faithful preclinical models^9–13^. Furthermore, the inability of conventional assays to capture cellular heterogeneity is particularly problematic when studying resistance, as it often arises from small subpopulations of cancer cells with distinct molecular properties.

To address these limitations, we applied the Teton assay that concurrently profiles RNA, protein, phospho-protein, and morphological features within individual cells^14^. Traditional multiomics approaches typically analyze different molecular layers using separate platforms, introducing variability and obscuring dynamic cellular responses. Recent advances in spatial and single-cell technologies highlight the importance of integrated, temporally resolved profiling for understanding complex cellular states^15–18^. Studies integrating RNA and protein data have provided more comprehensive insights into biological processes ^19^ and have revealed the rapid, transient nature of phosphorylation events, often occurring within an hour^20,21^. These dynamics are particularly relevant when studying therapeutic resistance, which frequently develops through clonal evolution, bypass signaling, or secondary mutations^22,23^. Our assay addresses these challenges by enabling the capture of molecular signatures across time points in a single experiment, offering a detailed view into drug-induced cellular responses.

As proof of concept, we applied this technology to NSCLC, where kinase inhibitors have shown promise but face resistance challenges. The targeted therapy era began with imatinib’s approval for BCR-ABL-driven leukemia, catalyzing kinase inhibitor development across multiple cancer types^24^. In NSCLC, EGFR inhibitors like gefitinib demonstrated remarkable efficacy in patients with EGFR-activating mutations^25–28^. However, resistance, often driven by the EGFR T790M mutation, typically emerges, leading to the development of third-generation inhibitors such as osimertinib^29^.

Combination therapies have emerged as a promising direction for overcoming resistance ^30–33^. In this study, we demonstrate how our integrative single-cell profiling approach reveals critical pathways involved in drug response and resistance. Through this molecular insight, we identified an effective combination therapy, which was independently validated in a previously published clinical study showing high efficacy^34^. These findings highlight the potential of single-cell multimodal analysis to generate actionable insights that translate into improved therapeutic strategies.

## Results

### TKI multiomic response in NSCLC

Gefitinib and osimertinib inhibit EGF receptor phosphorylation to reduce cell division and promote apoptosis (Figure 1A)^35,36^. We employed Teton assays to conduct a time-resolved study of how lung cancer cells respond to these drugs, monitoring MAPK/apoptosis and MAPK/cell cycle pathways (Figure 1B). We studied four NSCLC cell lines: A549 (wild-type EGFR, commonly studied for TKI sensitivity), NCI-H1299 (wild-type EGFR lacking p54 expression with a different genetic background), PC9 (EGFR exon 19 deletion mutant - most drug-sensitive), and NCI-H1975 (EGFR^T790M^ mutant conferring resistance to first and second generation TKIs)^36,37^.

**Figure 1.**
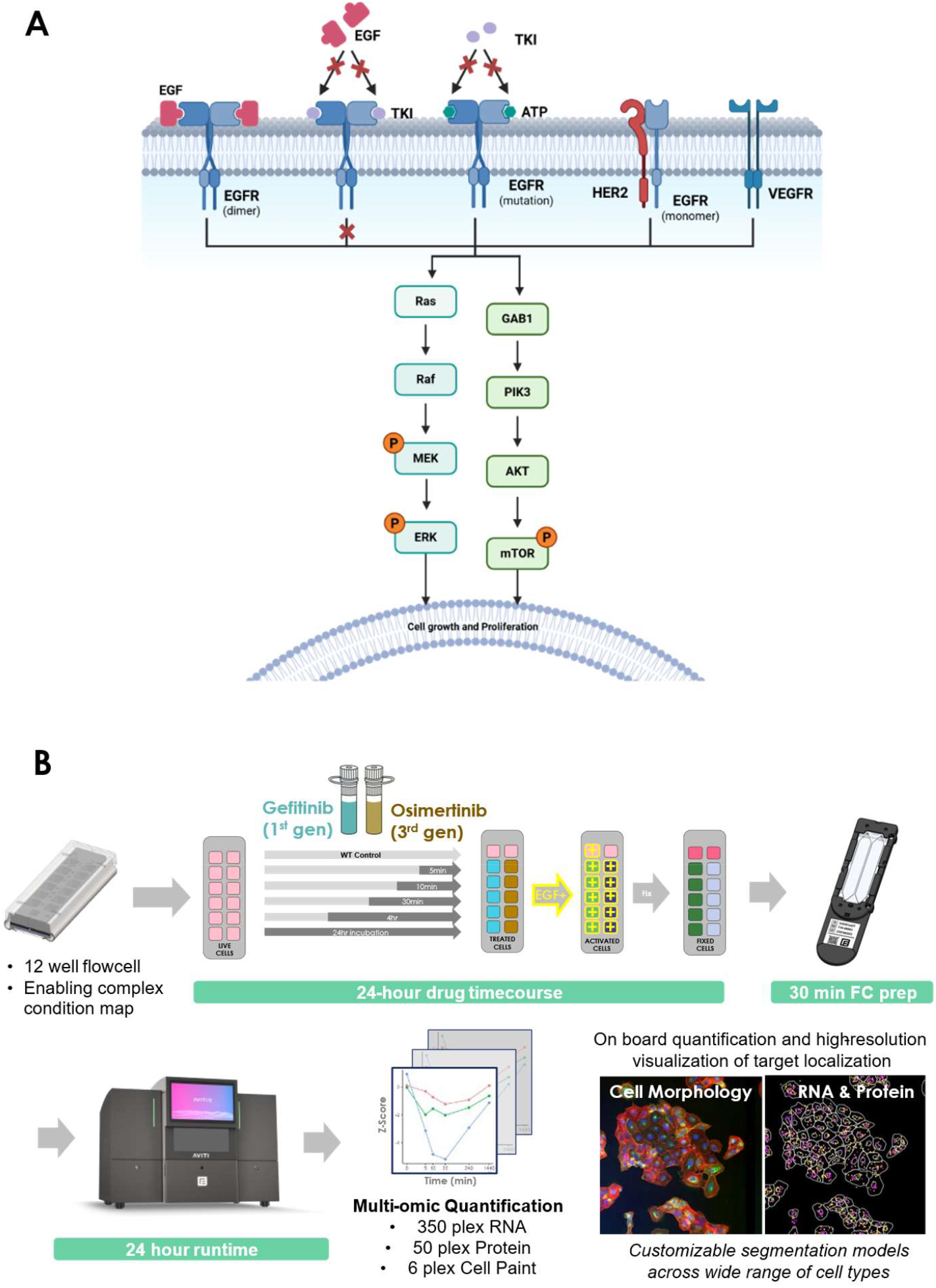
(A) Outline of mechanism of action for EGFR, TKIs, and established routes for resistance development in NSCLC cells, including expected impacted MAPK and related pathway. Figure 2 (A) Outline of mechanism of action for EGFR, TKIs, and established routes for resistance development in NSCLC cells, including expected impacted MAPK and related pathway. Red “X” marks represent blocking the binding site of EGFR domain using TKIs in wild-type EGFR, and ineffective inhibition and binding in mutant EGFR. Other escape pathways for TKI resistance include heterodimers or activation of alternative receptors. (B) Multiple NSCLC cell lines (A549, PC9, NCI-H1299, and NCI-H1975) were tested with the TKI drugs gefitinib and osimertinib in this study. Cells were seeded onto the Teton Slide Kit and time titrated with corresponding drugs, then assayed using a custom 55-plex protein panel and a 350-plex Cell Cycle RNA panel, with 6-plex cell paint.

At 100nM (lowest concentration), 30 minutes of exposure substantially reduced phospho-Y1068 EGFR in both A549 and PC9 cells, with higher concentrations nearly eliminating detection (Supplemental Figure 1). After 24 hours, all tested concentrations completely suppressed phospho-EGFR. Phospho-p44/42 ERK1/2 showed greater stability in A549 cells across all timepoints and concentrations, while in PC9 cells it was reduced at 10 μM after 30 minutes and at all concentrations after 24 hours.

Cell viability testing confirmed that while osimertinib was more potent than gefitinib, a 10μM concentration of either drug reduced cell populations in A549, NCI-H1299, and NCI-H1975 while still retaining over 50% of the original cell population, which was sufficient for multiomics analysis with the Teton assay (Supplemental Figure 2). This concentration was selected for subsequent experiments where cells were exposed to the drugs in a time course up to 24 hours, stimulated with EGF, and analyzed for differential responses across RNA (350 genes) and protein and phospho-protein (55 total proteins) focused on apoptosis, cell cycle, and MAPK/ERK pathway components, alongside morphological signatures using 6 cell paint markers (Supplemental Table 1).

**Figure 2.**
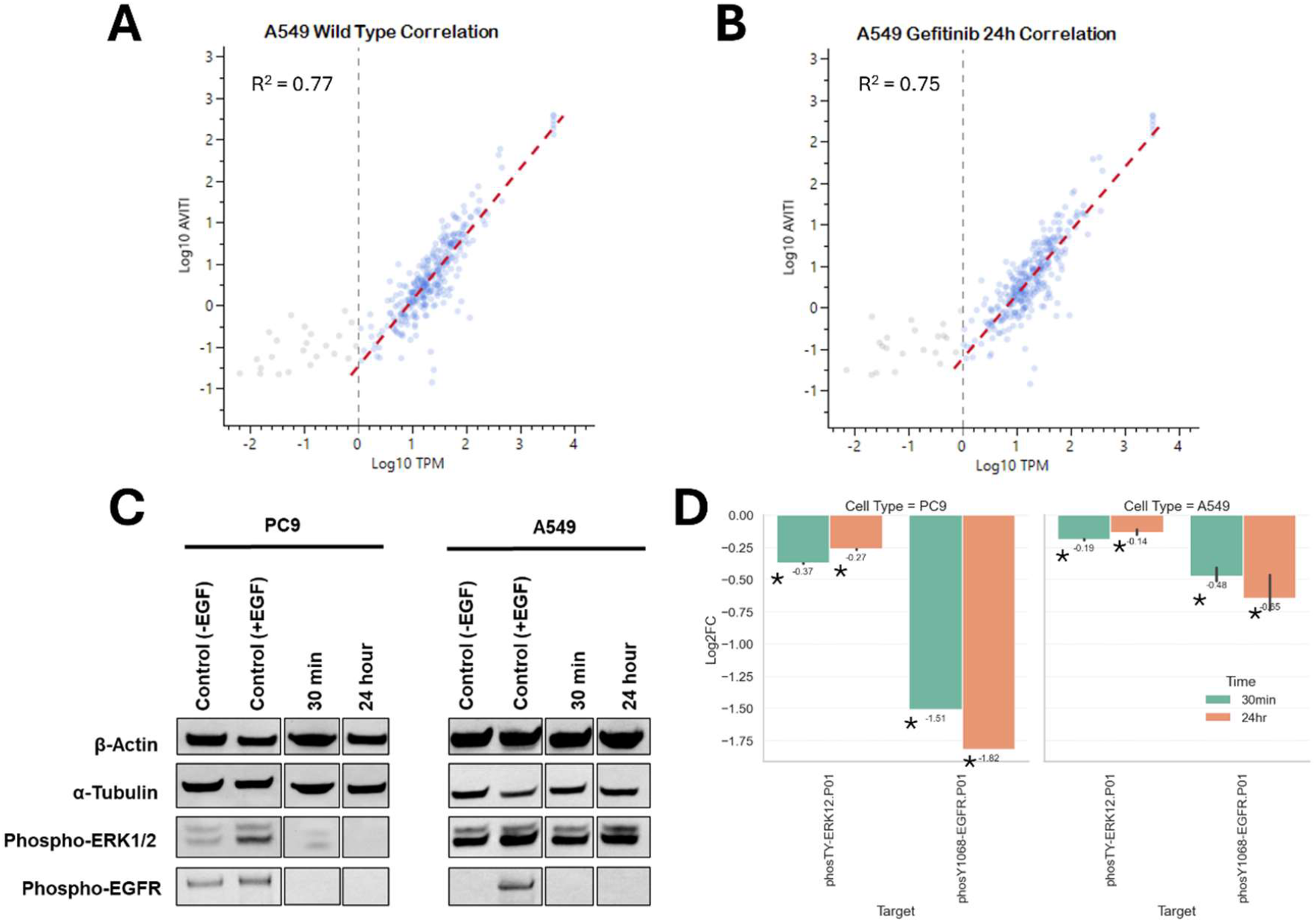
Validation of AVITI24 against established detection methods for RNA and protein. (A) Correlation of AVITI24 RNA counts and bulk RNA-seq of wild-type A549 cells. (B) Correlation of AVITI24 RNA counts and bulk RNA-seq of A549 cells treated with Gefitinib for 24hrs. (C) Comparison of western blot analysis of phospho-p44/42 ERK1/2 and phospho-EGFR in PC9 and A549 cells treated with 10uM Gefitinib for either 30min or 24 hours and (D) Log_2_ fold-change observed using Teton for phosphor-ERK1/2 and phosopho-ERK1/2 in PC9 and A549 cells after 30 minutes or 24 hours of 10uM Gefitinib treatment. The asterisk indicates significance at p < 0.05.

### Orthogonal Validation of Teton against standard RNA and protein detection methods

Building on previous validation across multiple cell types and cytokine responses^14^, we compared the Teton readout against established RNA and protein quantification methods, onfirming its applicability to NSCLC model systems. Parallel samples for bulk RNA sequencing (RNA-seq) and western blot were prepared. Comparison of the transcript abundance in both WT and treated cells showed high correlation (Figure 2A-B), with an apparent breakpoint in correlation at 1 TPM. Correlation was reported above this breakpoint.

To validate the Teton protein assay, we profiled phosphorylation of two canonical TKI response markers—phospho-Y1068 EGFR and phospho-p44/42 ERK1/2—in A549 and PC9 NSCLC cell lines treated with 10 μM gefitinib. These targets were selected based on prior studies showing EGFR down-regulation in both lines and differential ERK1/2 signaling, particularly minimal ERK response in A549 and sustained suppression in the more drug-sensitive PC9 line^35^.

Differential expression via the Teton assay was quantified as described in the Methods section. Using both western blot and Teton, we confirmed robust inhibition of phospho-Y1068 EGFR in both cell types. Phospho-p44/42 ERK1/2, as expected, remained largely unchanged in A549, but was suppressed in PC9 cells. Notably, Teton revealed additional temporal resolution: at 30 minutes, phospho-p44/42 ERK1/2 suppression was already detectable in PC9 (log_2_ fold-change = –0.37, p < 0.0001) and modestly reduced in A549 (log2 fold-change = −0.19, p < 0.0001). Phospho-Y1068 EGFR was significantly down-regulated at this early timepoint in both lines, with a stronger response in PC9, consistent with its enhanced sensitivity to EGFR inhibition.

By 24 hours, ERK1/2 phosphorylation largely recovered in A549 but remained partially suppressed in PC9, while EGFR phosphorylation remained low in both. While western blot captured these general trends, the Teton assay enabled more sensitive and quantitative detection of the dynamic changes, particularly at early time points, validating its utility for resolving subtle yet biologically meaningful shifts in signaling (Figure 2D).

### Bulk Population Response of TKI treated NSCLC cells

To better understand how EGFR mutational status influences downstream signaling dynamics, we compared MAPK/ERK pathway responses to EGFR TKI treatment across four NSCLC cell lines with varying EGFR genotypes (Supplemental Figure 3). A549 and NCI-H1299, both EGFR wild-type with different genotypes, were contrasted with the TKI-sensitive PC9 (harboring a single activating EGFR mutation) and NCI-H1975 (bearing both L858R and T790M EGFR mutations associated with resistance).

**Figure 3.**
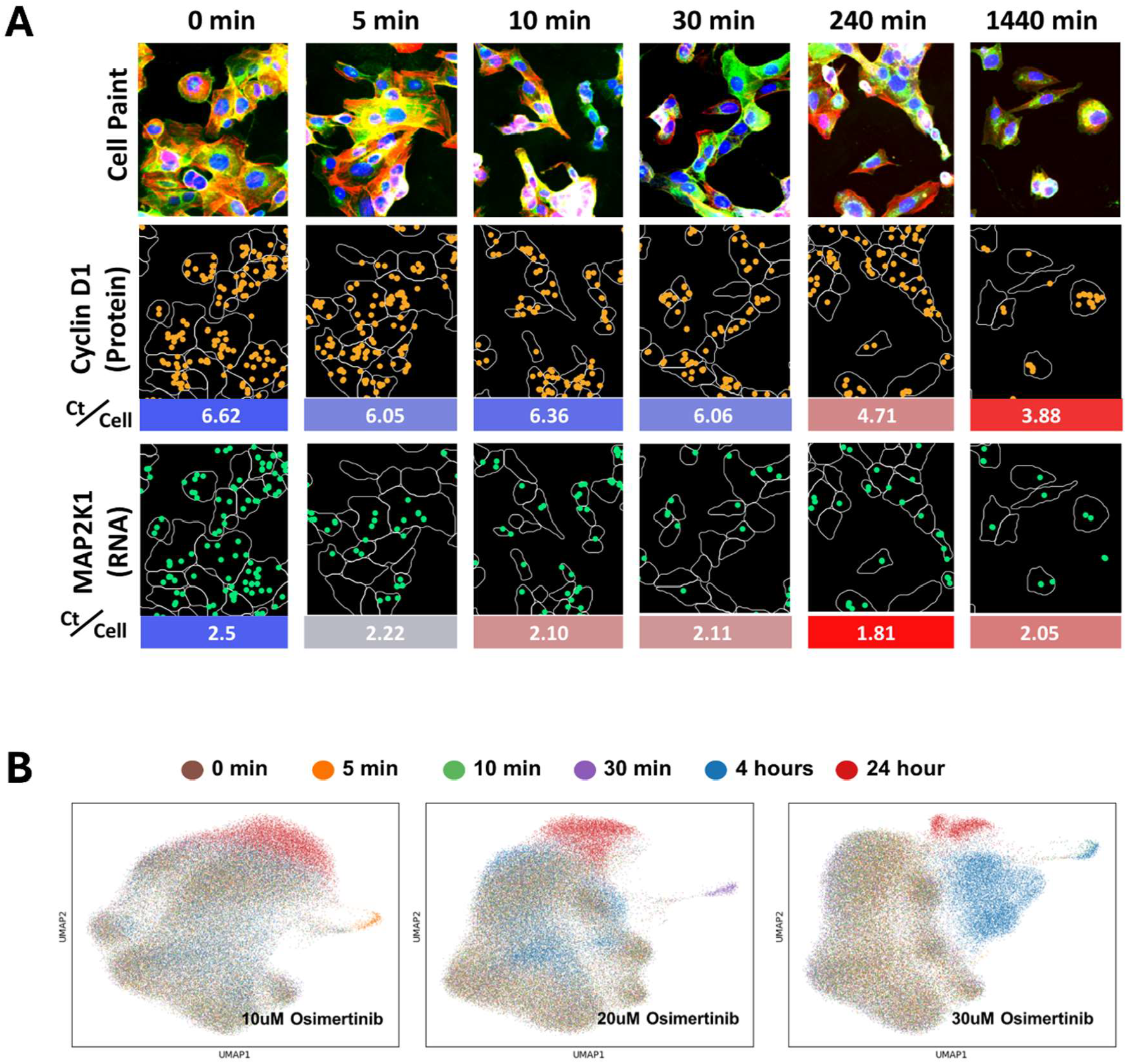
NCI-H1975 cells treated with 10uM Osimertinib for 0 to 24 hours. (A) Representative images for cell morphology and assigned protein and RNA counts. Bulk population averages for counts per cell (Ct/cell) are shown below targets cyclin D1 and MAP2K1. (B) UMAP projection of RNA response to TKI drug Osimertinib treatment at various concentrations for up to 24 hours. Each point represents a single cell detected within one AVITI24 run.

Upon Gefitinib treatment, PC9 cells exhibited a rapid and robust down-regulation of phospho-Y1068 EGFR as early as 5 minutes (log^2^ fold-change = −1.41, p < 0.0001), highlighting the immediate sensitivity of this genotype to first-generation EGFR inhibition. A549 cells demonstrated a reduced down-regulation response (log^2^ fold-change = −0.42, p < 0.0001). In contrast, NCI-H1299 and NCI-H1975 showed modest up-regulation after 5 minutes of exposure (log^2^ fold-changes = 0.40 and 0.12; p-values < 0.0001 and < 0..0001. respectively). These results suggest that depending on the genotype, 1^st^ generation TKI exposure may elicit a different response.

Osimertinib, a third-generation EGFR inhibitor, drove more immediate suppression across all lines. Notably, phospho-Y1068 EGFR was significantly down-regulated within 5 minutes in PC9, A549, NCI-H1299 and NCI-H1975, with effects maintained through 24 hours (log_2_ fold-changes: PC9 = −1.40, p < 0.0001; A549 = −0.70, p < 0.0001; NCI-H1299 = −0.22, p < 0.0001; NCI-H1975 = −0.25, p < 0.0001). The robust response across all cell lines and treatment times reflects the broader potency of Osimertinib regardless of EGFR mutation status.

Downstream in the pathway, phospho-p44/42 ERK1/2 levels were also significantly reduced by both TKIs. Gefitinib triggered measurable decreases across all four lines, with strong statistical support (Supplemental Figure 1), and osimertinib mirrored these effects with broad ERK suppression, confirming its activity across EGFR genotypes.

Interestingly, early transcriptional changes provided additional insight into pathway regulation. EGFR mRNA expression modestly increased at 30 minutes in PC9 after gefitinib treatment (log_2_ fold-change = 0.15, p = 0.014). Additionally, down-regulation was also observed in A549 cells after 24 hours of gefitinib treatment (log_2_ fold-change = −0.28, p = 0.043). Only modest and transient transcriptional response was observed in NCI-H1975 or NCI-H1299 with either TKI treatment, suggesting mutation-specific regulation of EGFR expression. In parallel, MAP2K2 expression rose in PC9 and A549 cells at 5 minutes but was transiently down-regulated in NCI-H1299 and NICI-H1975 following gefitinib exposure. Interestingly, in wild-type cells, MAP2K2 returned to baseline over time, suggesting a compensatory feedback mechanism that may act to re-engage MAPK signaling after initial suppression.

### Single-Cell Profiling Reveals Diverse and Cell-State-Specific Responses to TKI Treatment

To capture the heterogeneity underlying therapeutic resistance, we profiled NSCLC cells at single-cell resolution following EGFR TKI treatment, measuring RNA, protein, and morphological features over time. Since therapeutic resistance can emerge from rare subpopulations, this approach enabled a detailed dissection of drug responses that would be masked in bulk analyses.

Morphological changes were readily detectable following gefitinib and osimertinib exposure. Using a 6-plex Cell Paint targeting the membrane, nucleus, actin cytoskeleton, Golgi apparatus, mitochondria, and ER, clear changes in numerous morphological features supported RNA and protein results, while providing deeper insights into cell response (Supplemental Figure 6). Notably, at intermediate treatment times, cell area was reduced, while cells became more compact, an indicator of apoptosis^38^. Furthermore, mitochondria, endoplasmic reticulum, and nucleus intensity was decreased, suggesting compromised organelle integrity. Lastly, actin texture was reduced, while nuclear texture had increased, potentially indicating cytoskeletal disassembly and chromatin condensation. The severity of these changes was generally increased with osimertinib treatment, versus the 1^st^ generation gefitinib. At the molecular level, protein quantification revealed suppression of cyclin D1 (from 6.6 to 3.9 counts/cell) and MAP2K1 (from 2.5 to 1.8 counts/cell), reinforcing the drug’s impact at both phenotypic and signaling levels (Figure 3A, Supplemental Figure 1).

Transcriptional profiling revealed further complexity. UMAP projections of osimertinib-treated NCI-H1975 cells showed increasing stratification of RNA profiles with treatment time and dose. At 4 hours, divergence emerged at higher concentrations (20–30 μM), and by 24 hours, distinct subpopulations were present across all doses (Figure 3B). These results highlight a dose-dependent bifurcation in cell state in response to EGFR inhibition.

To resolve functional responses within the cell cycle, we trained a custom classifier to assign cells to G1, S, G2, or M phase based on RNA features (Methods). This enabled phase-specific analysis of TKI effects. CDCA3, a known prognostic biomarker, was significantly down-regulated only in G2-phase cells after 24 hours of gefitinib treatment in A549 (log_2_ fold-change = −0.47, p = 0.002). However, NCI-H1299 and NCI-H1975 showed significant down-regulation in G2 (log_2_ fold-change = −0.76, p = 0.003), M (log_2_ fold-change = −0.39, p = 0.049) and S (log_2_ fold-change = −0.23, p = 0.02) and G2 (log_2_ fold-change = −0.48, p = 0.019) and M (log_2_ fold-change = −0.67, p = 0.01) states, respectively (Figure 4A, Supplemental Table 2). Similarly, PLK1, an oncogenic kinase in NSCLC, showed phase-specific differential down-regulation in NCI-H1299 (G2 and S) after 24 hours of treatment, whereas NCI-H1975 and A549 showed down-regulation in all cell states (Figure 4B). These findings underscore the importance of resolving drug effects by cell state, as bulk-level averaging would obscure these phase-specific responses.

**Figure 4.**
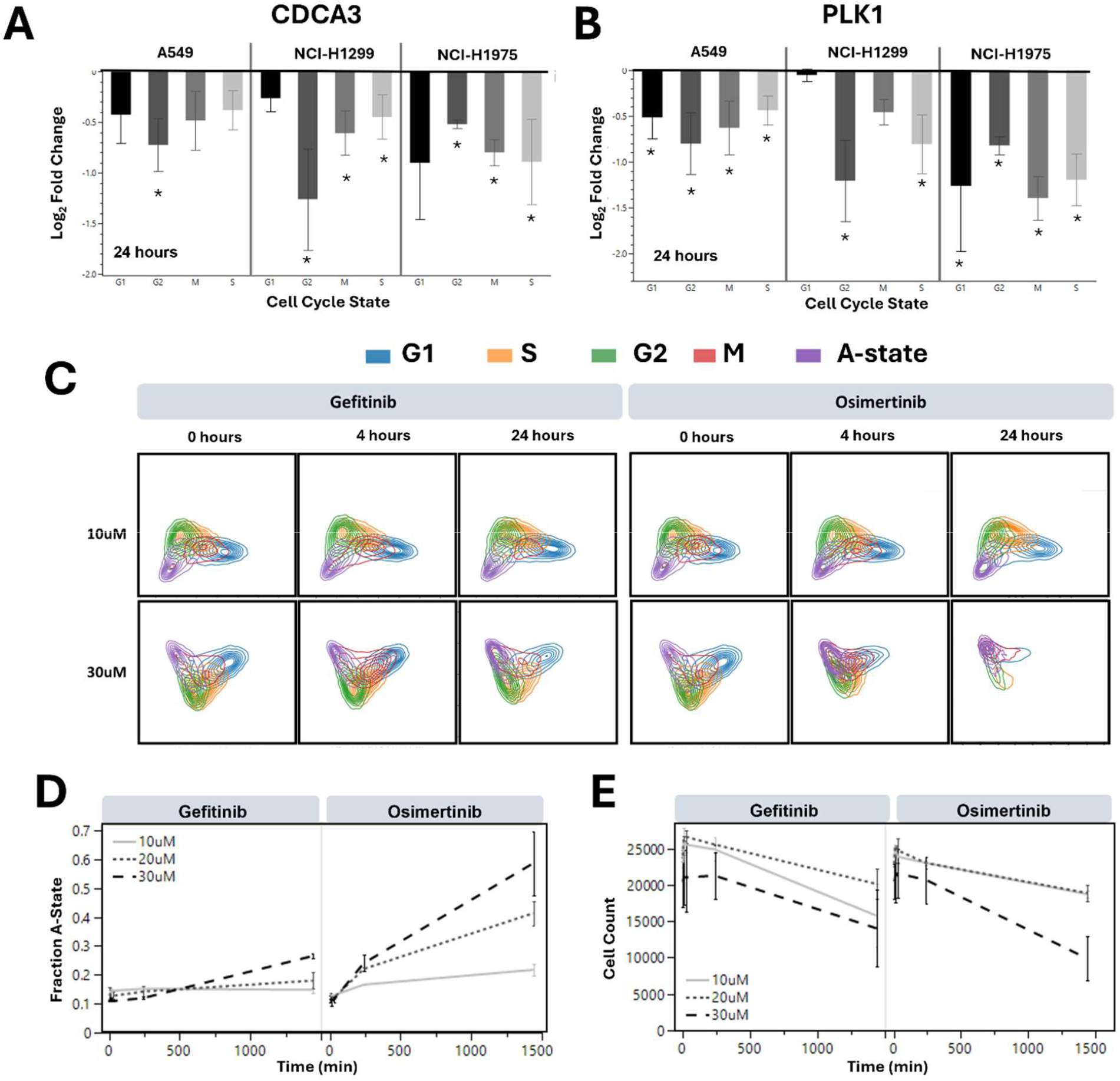
Cell population analysis of A549, NCI-H1299, and NCI-H1975 cells treatment with Gefitinib for 24 hours. (A) CDCA3 and (B) PLK1 RNA response stratified by cell cycle. (C) Factor analysis of cell populations for A549 cells treated with either Gefitinib or Osimertinib at 10uM and 30uM. (D) Fraction of cells in the A-state for A549 cells after treatment with Gefitinib and Osimertinib at variable concentrations and (E) quantified cell count. The asterisk indicates significance at p < 0.05.

Unexpectedly, the classifier also revealed a fifth, previously unannotated cell state (state A) that increased in abundance with treatment (Figure 4E). This population was defined by coordinated up-regulation of all APOP panel transcripts and exhibited a mixed transcriptional profile indicative of both pre-apoptotic signaling and survival-associated gene expression. The state was interpreted as a poised, apoptosis-primed but pro-survival phenotype, potentially reflecting a stress-induced equilibrium where autophagy and anti-apoptotic programs counterbalance pro-death signals, enabling cells to resist full apoptotic commitment.

Further analysis in A549 cells confirmed the relevance of this state to treatment outcome. High-dose osimertinib (30 μM) induced a pronounced increase in the fractional abundance of the fifth state compared to both gefitinib and lower-dose osimertinib. At 10 μM osimertinib, M-phase cells were selectively depleted by 24 hours, whereas at 30 μM, all cell cycle phases were affected (Figure 4C, Supplemental Table 2). Notably, the abundance of the fifth state increased by 79% and 169% under 30 μM gefitinib and osimertinib, respectively, compared to their 10 μM treatments (Figure 4D). This expansion correlated inversely with total cell counts (Figure 4G), suggesting that the cells remaining at higher drug doses disproportionately belonged to this survival-associated state. Rather than representing apoptotic escape, these cells may instead reflect a drug-tolerant population actively resisting death through enhanced survival signaling.

Together, these findings reveal a complex and adaptive landscape of single-cell responses to EGFR inhibition, ranging from early transcriptional divergence and cell cycle-specific effects to the emergence of a cell state potentially poised between death and resistance that may underlie residual disease and resistance.

### Targeting MAPK/ERK Signaling and Cyclin D1-CDK4 Axis Reveals Combination Vulnerabilities in NSCLC Cells

To dissect the impact of EGFR inhibition on MAPK/ERK and related signaling pathways, we profiled NCI-H1975 cells at the RNA, protein, and phospho-protein levels following 24-hour treatment with gefitinib or osimertinib (Figure 5). Both TKIs induced overexpression of EGFR mRNA, yet only gefitinib led to a rebound in phospho-Y1068 EGFR, suggesting differential feedback regulation at the receptor level (Supplementary Table 1). Across treatments, notable transcriptomic changes were detected for AKT1, RAF, and Rb, while protein-level suppression was observed in key signaling intermediates, including phospho-cyclin D1, cyclin D1, phospho-p44/42 ERK1/2, K-Ras, SOS1, and phospho-MEK1.

**Figure 5.**
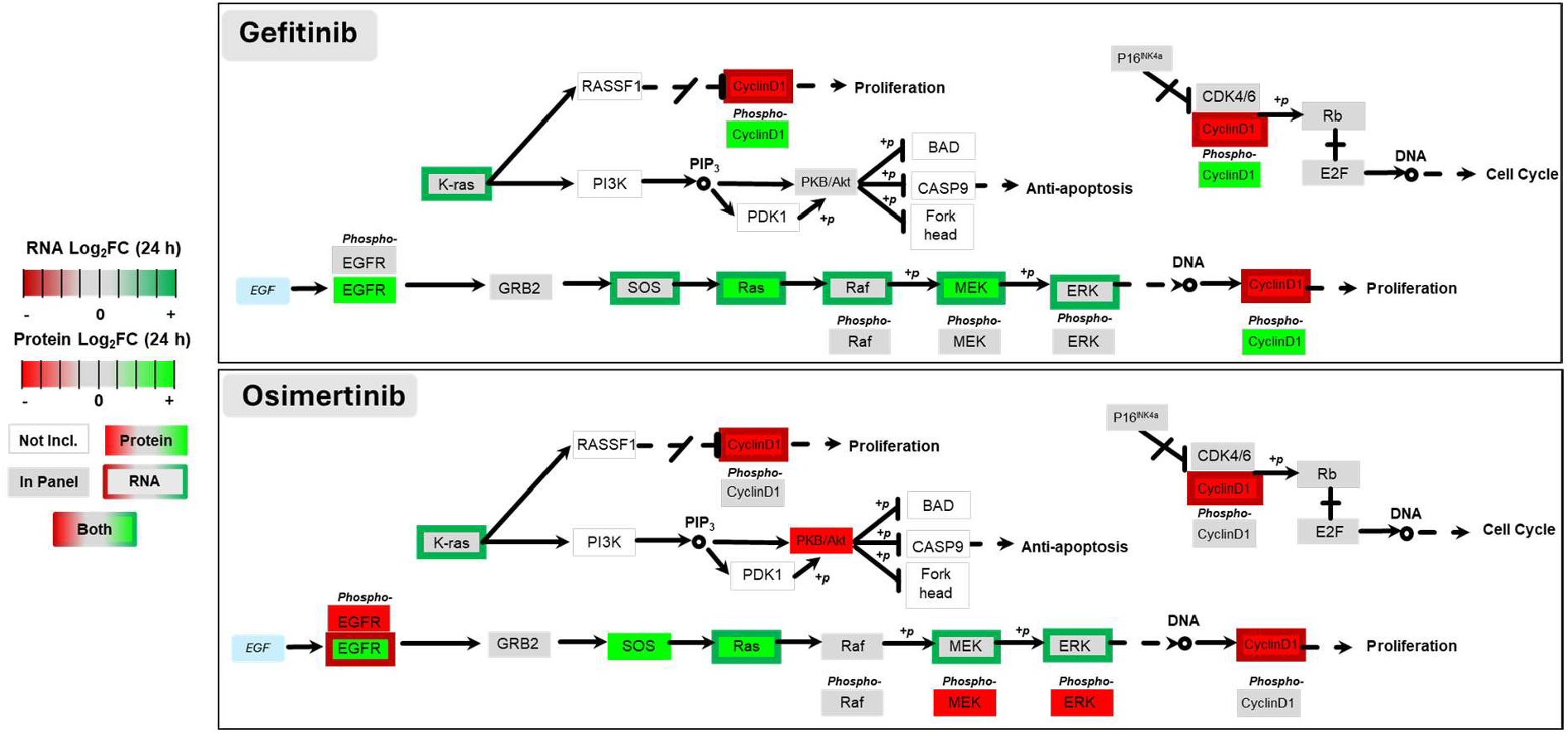
Pathway overview and summarized protein and RNA responses after 24-hour treatment with Gefitinib and Osimertinib at 10uM concentration on NCI-H1975 cells. Gray blocks include targets within Teton cell cycle panel, fully colored blocks are regulated proteins, colored outlined blocks are regulated RNA targets, white blocks are not included with the panels.

Temporal profiling revealed a muted response at early time points: gefitinib produced limited suppression of phospho-proteins at 4 hours, and MEK phosphorylation remained only partially inhibited throughout. Interestingly, cyclin D1 protein levels were selectively reduced by osimertinib, suggesting a post-transcriptional block in cell cycle progression uniquely triggered by third-generation EGFR inhibition.

Given the role of cyclin D1-CDK4 interaction known to drive cell survival, we next evaluated whether co-targeting this axis could enhance the efficacy of TKI treatment. Using the CDK4 inhibitor palbociclib in combination with either gefitinib or osimertinib, we observed further modulation of cyclin D1 signaling in A549 cells. Compared to TKI treatment alone, the combination therapy resulted in modestly elevated levels of both cyclin D1 and phospho-cyclin D1 (log2 fold-changes = 0.32 and 0.15; p < 0.0001 and p = 0.0004 with 1 μM palbociclib), respectively for gefitinib (Figure 6C). Combination treatment with osimertinib did not show significant changes. This upregulation, paradoxical at first glance, likely reflects a compensatory feedback response driven by inhibition of downstream targets.

**Figure 6.**
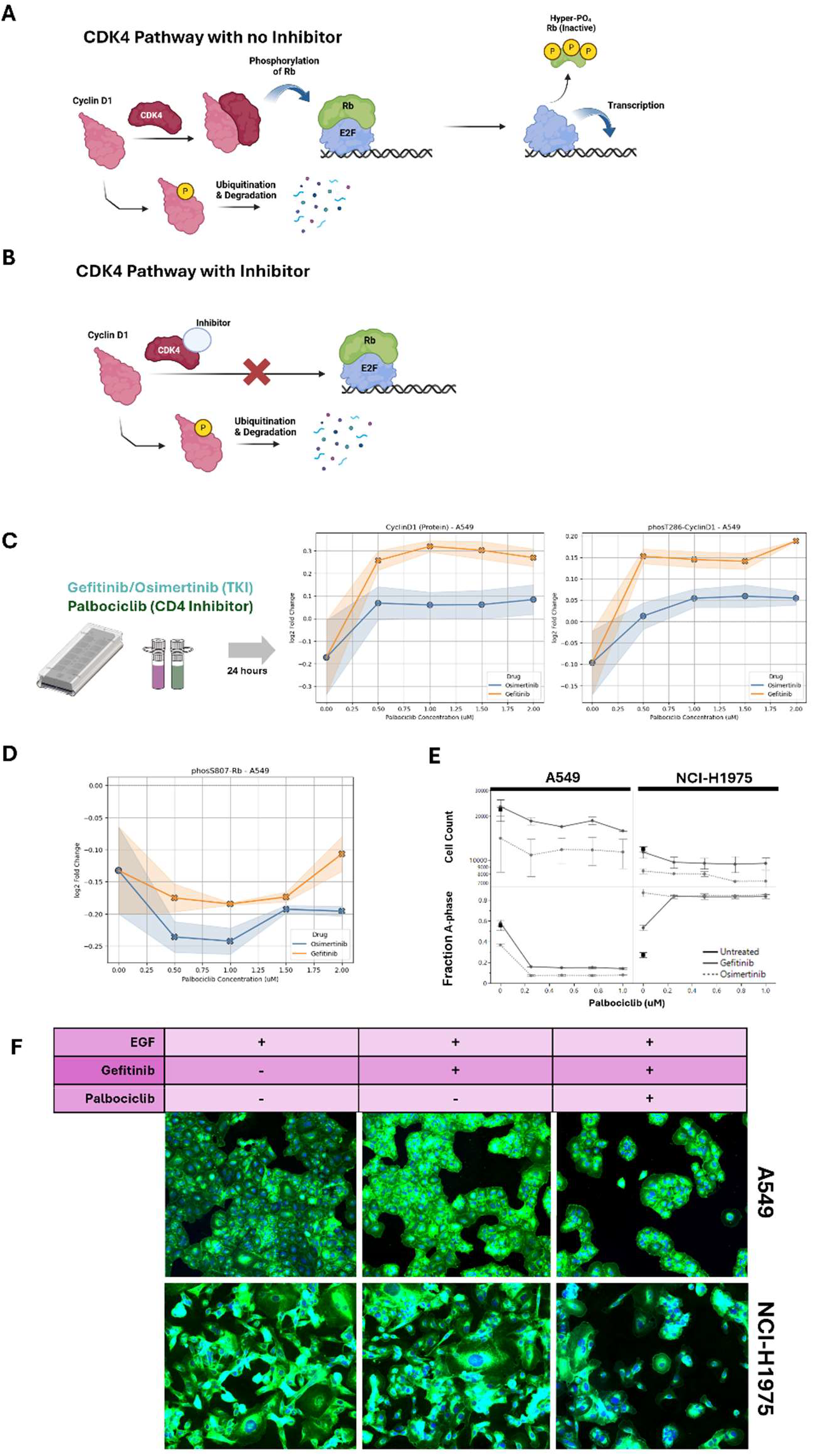
(A) CDK4 pathway for cell survival via induced transcription with the E2F transcription factor. (B) CDK4 pathway arrest with CDK4 inhibitor to prevent release of E2F and promotion of cyclin D1 degradation. (C) Workflow for combinatorial drug treatment with 20uM TKI and CDK4/6 inhibitor palbociclib for 24 hours, to combat potential cell survival, and cyclin D1/phospho-cyclin D1 response to drug combination. (D) Response for downstream Phos-Rb (protein). (E) Cell count and A-state cell fraction for A549 and NCI-H1975 with gefitinib/palbociclib or osimertinib/palbociclib treatment. (F) Representative images for A549 and NCI-H1975 response to gefitinib and palbociclib.

Despite this, downstream markers of CDK4 activity were consistently suppressed with combination therapy. Notably, phospho-Rb (S807), a key CDK4 substrate, was reduced in both conditions (e.g. at 1 μM of palbociclib - gefitinib: log_2_ fold-change = −0.18, p < 0.0001; osimertinib: −0.24, p = 0.0001), confirming effective disruption of the cyclin D1-CDK4-Rb axis (Figure 6D).

These molecular effects translated into dramatic reductions in cell viability (Figure 6E). In A549, gefitinib/palbociclib and osimertinib/palbociclib combinations decreased viable cell counts by 79% and 80%, respectively. Notably, NCI-H1975 cells, which are more resistant to monotherapy, were even more sensitive to the combination treatment, with reductions of 86% and 96%, respectively. These effects were supported by decreased cell confluency (Figure 6E–F), underscoring the cytotoxic potential of dual pathway blockade.

Finally, we examined whether the emergent A state correlated with cell viability. In A549 cells, a strong inverse relationship was observed: increased abundance of the A state tracked with greater cell loss, suggesting it marks a drug-sensitive or apoptosis-primed population. In contrast, this association was absent in NCI-H1975, implying that other resistance mechanisms may be present in this context.

Together, these results highlight the value of targeting cell cycle machinery in conjunction with EGFR inhibition and point to cell state-specific vulnerabilities that may inform future therapeutic strategies.

## Discussion

In this study, we utilized the AVITI24 multiomics platform to dissect cellular signaling in NSCLC during TKI exposure, demonstrating its capacity to accelerate scientific discovery in this clinically important setting. Current approaches to study treatment resistance rely on established methods that analyze cellular responses through separate experiments for different molecular readouts, typically measuring a restricted set of targets at once. Our technology overcomes these limitations by simultaneously capturing RNA, protein, phosphoprotein, and morphology in single cells to enhance understanding of tyrosine kinase inhibitor mechanisms in both wild-type and mutant EGFR NSCLC cells.

The Teton assay demonstrated high correlation with traditional methods while offering several distinct advantages. When compared to western blotting, it provided superior sensitivity and enabled subpopulation identification not possible with bulk RNA sequencing. The platform’s ability to simultaneously assess a panel of proteins alongside transcriptional changes streamlines the experimental workflow compared to RNA sequencing protocols, eliminating separate library preparation steps while maintaining comparable data quality.

While our approach demonstrates promising results, several technical and analytical limitations should be acknowledged. First, although we collected a wealth of multimodal data, this proof-of-concept study presents only a fraction of possible analyses, leaving substantial opportunity for deeper exploration. Additionally, because the morphological responses to treatment were largely exploratory, some of these findings remain open to interpretation and warrant further validation. Our protein panel, though informative, was limited to 55 targets, and future expansions will enable broader coverage of relevant pathways. Finally, detection limits for both protein and RNA constrained us to focus on relative expression changes, rather than absolute quantification of low-abundance molecules.

Despite these limitations, single-cell resolution enabled us to uncover heterogeneity in drug response that would remain hidden in bulk population analyses. When examining osimertinib titrations, we identified responsive subpopulations across a range of drug concentrations, offering insights into the spectrum of cellular responses. Applying a cell cycle model to stratify NSCLC cells by phase revealed distinct phase-specific sensitivities, providing mechanistic clues to drug action.

The superior efficacy of osimertinib over gefitinib for NSCLC was evident throughout our study, particularly in TKI-resistant cell lines where gefitinib’s binding is compromised by steric mutations. Osimertinib’s covalent binding to T790M mutant EGFR enables more effective treatment of resistant lines like NCI-H1975. A key advantage of our approach was the ability to monitor pathway responses at very early timepoints after drug exposure. We observed that these first-generation and third-generation TKIs elicited similar responses in upstream EGFR signaling, but diverged at the level of MEK and ERK, particularly in resistant cell lines. This temporal resolution allowed us to capture the rapid phosphorylation dynamics of the MAPK pathway and identify points where signaling persisted despite TKI treatment.

Our most significant finding emerged from this pathway analysis, where we discovered that cyclin D1 expression was not sufficiently downregulated with gefitinib treatment compared to osimertinib, indicating a potential bypass mechanism for cellular survival. The cyclin D1-CDK4 heterodimer facilitates retinoblastoma protein phosphorylation, releasing E2F transcription factors that promote cell cycle progression. By combining CDK4 inhibitor palbociclib, shown previously to have only modest effects as a monotherapy in NSCLC^39^, with either TKI, we significantly enhanced therapeutic efficacy. This combination improved the potency of first-generation gefitinib and further increased third-generation osimertinib efficacy in both wild-type EGFR A549 cells and double-mutant NCI-H1975 cells. The clinical relevance of this TKI-CDK4 inhibitor combination has been independently validated in clinical settings^34,40^ affirming the translational value of insights obtained through our study.

## Conclusion

Our study demonstrates the power of integrated multiomic analysis at single-cell resolution to uncover mechanisms of drug resistance in NSCLC. By simultaneously measuring RNA expression, protein levels, phosphorylation states, and cellular morphology in the same cells, we identified cell cycle-dependent responses to TKIs and discovered that cyclin D1-CDK4 activation provides an escape route from TKI-induced cell death. This insight led to a targeted combination therapy strategy using palbociclib with either gefitinib or osimertinib, significantly enhancing treatment efficacy even in EGFR-mutant cells that typically resist first-generation TKIs.

This work highlights how single-cell multiomic profiling can reveal heterogeneous responses within seemingly homogeneous cell populations, identifying resistant subpopulations that would be masked in bulk analysis. Our demonstration that a first-generation TKI can achieve efficacy comparable to a third-generation TKI when appropriately combined with targeted CDK4 inhibition offers a promising therapeutic strategy that aligns with independent clinical studies. Beyond NSCLC, this approach provides a framework for investigating complex drug resistance mechanisms across diverse cancer types, potentially accelerating the development of precision combination therapies.

## Methods

### Instrumentation

Element Biosciences multiomics platform was leveraged through the AVITI24 (880-00004, Element Biosciences, San Diego, CA) instrument, employing the Teton Human MAPK & Cell Cycle Kit, 12 Well (860-00023), Teton Human MAPK & Apoptosis Kit, 12 Well (860-00025), Teton, Flow Cell Assembly Kit, 12 Well (2-pack, 860-00028), Teton Slide Kit, Uncoated - 12 Well (2-pack, 860-00032), and Teton Flow Cell Assembly Tool Set (860-00033).

Fluorescent microscope (Olympus IX83) equipped with Z drift component (IXZP-ZDC2-830), Sutter Instrument Lambda 10-B Smart Shutter, ASI MS-2000 Flat-Top XY Automated Stage, Andor Zyla sCMOS 4.2P USP3.0 camera, filters for green channels (515–560 nm excitation, 580–650 nm emission) and red channels (620–650 nm excitation, 660–750 nm emission) and Olympus UCPLFLN20X fluorescence objective. General cell segmentation models were used for A549, NCI-H1299, and NCI-H1975. Hela cells were segmented using the Hela segmentation model.

### Cell Lines

Cell lines including A549 (ATCC CCL-185), NCI-H1299 (ATCC CRL-5803), and NCI-H1975 (ATCC CRL-5908) were purchased from the American Type Culture Collection (ATCC). A549 was cultured in F12K Medium (ATCC 30-2004) supplemented with 10% FBS and 1% Penicillin / Streptomycin (10,000U/mL stock). NCI-H1299 and NCI-H1975 were cultured in RPMI-1640 Medium (ATCC 30-2001) supplemented with 10% FBS and 1% Penicillin/Streptomycin (10,000U/mL stock, Gibco, 15140-122). All cell lines were grown at 37°C and 5% CO_2_ in a humidified environment.

### Cell Viability Assays

Cells were seeded in a 96-well plate at a density of 5000-8000 cells/well (depending on treatment duration and cell line) and allowed to adhere overnight. Cells were then exposed to various concentrations of kinase inhibitors gefitinib (MedChemExpress HY-50895), osimertinib (MedChemExpress HY-155772), and dacomitinib (MedChemExpress HY-13272), which were stored in DMSO and diluted such that the final DMSO concentration in the media was no more than 0.5% v/v. After 24 to 48 hours, cells were washed twice with 150uL per well warmed DPBS and fixed in 100uL per well 3.7% formaldehyde (Sigma-Aldrich, F8775-25mL) for 30min. Cells were then washed thrice with 150uL per well 1x PBS, permeabilized in 70% EtOH for 10min at RT, then incubated in 80uL per well of a mixture of 0.067X CellBrite® Orange (Biotium #30022) and 0.5X RedDot™2 (Biotium #40061-1) in PBS for 1hr at 37°C. Cells were then imaged on the fluorescent microscope to obtain cell counts for viability calculations. Percent cell viability was calculated relative to DMSO vehicle controls.

### Western blot analysis of NSCLC cells

Cells were seeded at appropriate densities in T75 flasks (9×10^5^ cells per flask for PC9, 1.5×10^6^ cells per flask for A549). At times required for the treatment timepoint (e.g. 24 hours prior to harvesting) media was replaced with pre-warmed media containing the appropriate concentration of gefitinib. Prior to harvesting the necessary flasks were stimulated with 200ng/mL EGF for 15 minutes, during which time gefitinib was still present in the media. Media was removed and retained, cells were dissociated using 0.25% Trypsin-EDTA (Gibco 25200-056), then the retained media was used to quench the Trypsin. A small aliquot of cells was saved for counting, then cells were centrifuged at 200rcf for 5 minutes, resuspended in 600uL TRI Reagent, and stored at −80°C. 400uL of each sample was processed downstream to extract protein for western blotting. 400uL of 100% ethanol was added to each sample followed by 3.2mL of prechilled acetone. Samples were chilled on ice for 30 minutes to allow the protein to precipitate. Then samples were centrifuged at 4000rpm for 1 minute. Supernatant was discarded, pelleted samples were rinsed with another 400uL of 100% ethanol and centrifuged at 2000rpm for 15 seconds. Supernatant was discarded again, and each pellet was resuspended in a volume of UltraPure Distilled Water (Invitrogen 10977-015) to approximate 7500 cells/uL. An ultrasonic processor wand (Sonics Vibra Cell) was used in 5 pulses of 5 seconds each to fully resuspend the pellets. A Bradford assay was conducted using Coomassie Plus Reagent (ThermoScientific 23238) to measure each sample’s protein content for normalization. Samples were diluted to 10ng using ultrapure water, 4x NuPAGE LDS Sample Buffer (Invitrogen NP0007) diluted to 0.6x, and 10x NuPAGE Sample Reducing Agent (Invitrogen NP0009) diluted to 1x for a final volume of 20uL. Samples were loaded into a 1.0mm x 12-well 4-12% Bis-Tris NuPAGE Gel (invitrogen NP0322BOX) alongside 5uL PageRuler Prestained Protein Ladder (ThermoScientific 26616) and run in fresh 1x MES SDS Running Buffer (Invitrogen NP0002) for 1 hour at 180V. Gels were then transferred to PVDF membranes using iBlot 2 (Invitrogen) and iBlot 2 PVDF Regular Transfer Stacks (Invitrogen IB24001). iBlot stack was separated to place gel smoothly on PVDF membrane, then filter paper was soaked in distilled water and placed on gel, then copper top stack was placed on filter paper, followed by absorbent pad on top stack. iBlot 2 was run for 7 minutes: 20V for 1 min, 23V for 4 min, 25V for 2 min.

After transferring, stacks were disassembled, and membranes were transferred to sterile petri dishes. Membranes were first rinsed in distilled water then submerged in EveryBlot Blocking Buffer (BioRad 12010020) for 17.5 hours (overnight). For all incubation and wash steps membranes were kept in sterile petri dishes on a shaker with gentle agitation in a room at 4C. Blocker was drained and membranes were then incubated with primary antibodies plus control antibodies diluted in Everyblot as such: Anti-EGFR (phospho Y1068) antibody (Abcam AB40815) was diluted 1:500 for a final concentration of 1.908ug/mL along with control β-Actin antibody (ProteinTech 81115-1-RR) diluted 1:10,000 to 0.1ug/mL. Phospho-p44/42 MAPK (ERK1/2) antibody (Cell Signaling Technology 4370S) was diluted 1:1000 to 0.502ug/mL along with control α-Tubulin (Abcam AB52866) diluted 1:20,000 to 0.324ug/mL. Membranes were incubated with primary antibodies for 2 hours, then washed 5 times for 5 minutes each using Tris buffered saline + 0.1% Tween 20 (TBST). Membranes were incubated in Secondary antibody Goat Anti-Rabbit IgG H&L (HRP) (Abcam AB6721) diluted 1:2500 to 0.8ug/mL in EveryBlot plus 0.02% SDS for 2.5 hours, then washed 6 times for 5 minutes each using TBST. For fluorescent imaging membranes were incubated in a 1:1 ratio of SuperSignal West Dura Extended Duration Substrate (ThermoScientific 34075) reagents for 5 minutes with agitaiton and then immediately placed on a Typhoon FLA 9500 (GE Healthcare Life Sciences) using a Cy2 method with 500V PMT and 50uM pixel size.

### Well level cell population RNA Sequencing

RNA of the A549 cell line was isolated and purified using the Direct-Zol RNA Miniprep (Zymo Research R2053) kit and protocol. Samples in 80μL TRI Reagent® were thawed and 80μL 100% ethanol was added. Mixtures were vortexed and transferred to the provided spin columns and centrifuged. All centrifugation steps were performed at 10,000 x g for 30 seconds. After discarding the flow through, 400μL RNA Wash Buffer was added to the column and centrifuged. DNase I was diluted 1:16 in DNA Digestion Buffer to a final concentration of 0.375 U/μL in 80uL, added to the column, and incubated for 15 minutes at room temperature. Then columns were washed twice with 400μL of Direct-zol RNA PreWash and once with 700μL of RNA Wash Buffer. Final purified RNA was eluted in 50μL of DNase/RNase-Free water and stored in RNase-free tubes at −80°C until bulk RNA sequencing.

Following RNA extraction, the RNA Library Prep Kit (Watchmaker, 7K0078-096) and RNA Library Prep Kit with Polaris™ Depletion (Watchmaker, 7BK0002-096) were utilized according to standard library preparation protocols, then sequenced using AVITI 2×150 Elevate Sequencing Kit.

### Custom antibody add-on screen for Teton protein panel

Cells were seeded in a 96-well plate at a density of 5000-8000 cells / well and allowed to adhere overnight, then cells were washed twice with 150uL per well warmed DPBS and fixed in 100uL per well 3.7% formaldehyde (Sigma-Aldrich, F8775-25mL) for 30min. Cells were then washed thrice with 150uL per well 1x PBS, permeabilized in 70% EtOH for 10min at RT, then incubated in 80uL per well of primary antibody diluted according to manufacturer recommendations in 1X PBS to perform immunofluorescence. After an hour incubation at room temperature, cells were washed thrice with 150uL 1X PBS, then incubated with 80uL of Alexa Fluor® 647 Donkey anti-rabbit IgG (Biolegend #406414) diluted to manufacturer’s recommendation in 1X PBS for 1 hour at room temperature. Cells were washed thrice with 150uL 1X PBS, then another 150uL 1X PBS added to the wells. The cells were imaged on the fluorescent microscope to observe immunofluorescence competency.

Hits that demonstrated sufficient signal via immunofluorescence that corresponded to expected phenotypes were tested via Teton Antibody Screening Kit to further down-select targets with sufficient counts/cell to add to our custom protein panel. Primary antibodies for phosT286-cyclin D1 (CST #3300S), phospho-Y1068 EGFR (Abcam ab40815), Ras (Abcam ab52939), SOS1 (Abcam ab140621), cyclin D1 (Abcam ab134175), and EGFR (Abcam ab52894) were used as custom additions via the Teton Custom Add-On Protein Panel Assembly Kit.

### Time-dependent cellular response to kinase inhibition

Cells were seeded in an Element Biosciences Teton™ CytoProfiling flow cell coated with Poly-L-lysine (Sigma P4707) according to manufacturer specifications at 7,000 cells / well for A549, 7,000 cells / well for NCI-H1299, 10,000 cells / well for NCI-H1975, and 7,000 cells / well for PC9, then allowed to adhere overnight. At 24 hour, 4 hour, 30 minute, 10minute, and 5minute pre-EGF treatment, existing media was removed from the well and replaced with media containing 10μM gefitinib or 10μM Osimertinib to a final DMSO vehicle concentration not exceeding 0.5% v/v. After the timepoint durations were complete, media was removed from wells and replaced with media containing 10μM gefitinib or 10μM Osimertinib (respective to the treatment) and 200ng/mL EGF (Sigma E4127). Cells were incubated at 37°C for 15min in this media, then washed twice with 200uL per well of warmed DPBS and fixed in 150uL per well of 3.7% formaldehyde (Sigma-Aldrich, F8775-25mL) for 20min, then washed twice with 200uL per well 1x PBS then stored at 4°C in PBS containing 0.1U/uL RiboLock RNase Inhibitor (Thermo Scientific EO0381) until use on AVITI24 or prepared for bulk RNAseq by washing twice with 200uL per well warmed DPBS then lysed in 100μL per well TRI Reagent® and stored at −80°C until prepared for sequencing. On AVITI24, flowcells were sequenced using the Teton Human MAPK & Cell Cycle Panel Kit (Element Biosciences, 830-00040), Teton Human MAPK & Apoptosis Panel Kit (Element Biosciences, 830-00041) according to user guidance protocols.

### AVITI24 differential expression analysis

Differential expression from the Teton assay was quantified by comparing the log-ratio of each target’s abundance relative to the control condition. It was assumed most targets in the comparison were invariant, and the standard deviation of the null distribution was estimated as 1.48 times the median absolute deviation of the log-ratios. A Z-score was calculated as the log ratio divided by the standard deviation to quantify deviation from the control condition of the cell line with EGF activation and no TKI treatment. P-values were calculated from the standard normal distribution and used to determine significant changes in expression.

### Cell cycle state modeling on AVITI24

Hela cells (ATCC, CCL-2) were seeded onto PLL coated Teton™ CytoProfiling 12 well flow cell at 4500 cells per well in culture media (DMEM (Gibco, 10566-016) plus 10% FBS (Cytiva, SH30396.03HI), 1% PenStrep (Cytiva, SV30010), 1% MEM NEAAs (Gibco, 11140,050), and 1% Sodium Pyruvate (Gibco, 11360-070)) and incubated overnight to allow cells to attach to surface. Media was then replaced with media with cell phase arresting agents for: S (Thymidine, 2mM, Sigma T1895-1G), G2 (RO-3306, 10uM, Sigma 217699-5mg), M (Nocodazole, 0.2ug/ml, Stemcell Technologies 74072), G1 (Lovastatin, 20uM, Sigma 438185-25mg), G1/S (Aphidicolin, 5ug/ml Sigma 178273-1mg) and allowed to incubate at 37°C for 20 hours before fixation. Cells were washed twice with 200uL per well of warmed DPBS and fixed in 150uL per well of 3.7% formaldehyde (Sigma-Aldrich, F8775-25mL) for 20min, then washed twice with 200uL per well 1x PBS then stored at 4°C in PBS containing 0.1U/uL RiboLock RNase Inhibitor (Thermo Scientific EO0381) until used on AVITI24. Flowcell was run on AVITI24 using the Teton Human MAPK & Cell Cycle Kit (Element Biosciences, 860-00023). Two replicate flowcells were run on AVITI24.

Following run completion, transcript counts for both AVITI24 runs were normalized per cell by total detected counts per batch and outlier cells were excluded via cell area and assigned counts. Cells were then combined across both runs for further analysis. Further preprocessing normalized cells across all sequencing batches and log-transformed counts. Transcript features were projected into their principal components. Upon inspection of the first two principal components, a two-dimensional annulus was observed, consistent with prior observation^41^ (Supplemental Figure 4). Cells from wells with different synchronization agent treatments were observed to cluster in different regions of the annulus. Unsupervised clustering was performed on the PCAs using the Leiden algorithm to achieve the expected number of clusters^42^. Following clustering, clusters were assigned to each cell cycle state based on their position on the annulus and their enrichment of cells synchronized in a specific state. This resulted in four clusters for each cell cycle state. Then an enrichment analysis was performed for each state relative to the others using a t-test method. This resulted in an enrichment score for each transcript in each cluster relative to the others. The top 20 transcripts for each state were then used in other AVITI24 runs to score cells for the most likely cell cycle state via methods described previously^43^.

### A-state classification and interpretation

The A-state classification was performed using up-regulation all Teton APOP-panel RNA targets for each experiment relative to the four cell cycle states as defined above methods described previously^43^. Cell count pre-processing was performed as was done for cell cycle classification. Interpretation of the A-state was performed by calculating the log_2_ fold-change and Wilcoxon rank-sum test for each RNA target relative to other cells. Robustly up and down regulated targets were determined by taking the minimum test statistic across a total of 12 Teton runs (Supplemental Figure 5).

### Morphology Analysis

Morphological profiling was performed using all six Cell Painting channels and corresponding segmentation masks using CellProfiler^44^. The AreaShape module was used to compute geometric properties of each cell based on the segmented boundaries. Per-cell intensities were calculated by subtracting background and summing pixel values within the segmented cell area for each Cell Paint feature. Additional morphological features were extracted using the MeasureObjectSizeShape, MeasureGranularity, MeasureObjectIntensity, MeasureObjectIntensityDistribution, and MeasureTexture CellProfiler modules. In total, this pipeline yielded 575 features per cell, spanning intensity, texture, granularity, and shape descriptors across all six imaging channels.

To quantify morphological shifts between treated and control conditions, we computed a signed, per-feature Kolmogorov–Smirnov (KS) statistic across matched replicate wells. For each treatment, wells were paired with control wells in sorted order. For every feature and well pair, we computed a signed KS statistic reflecting both the magnitude and direction of distributional shifts.

### Combinatorial therapy with gefitinib/osimertinib and palbociclib

7,000 cells per well A549 and 10,000 cells per well NCI-H1975 were seeded in an Element Biosciences Teton™ CytoProfiling flow cell coated with Poly-L-lysine. Cells were allowed to adhere overnight and then challenged with a mix of gefitinib or osimertinib combined with palbociclib (PD0332991, MedChemExpress HY-50767) for 24 hours. Cells were then stimulated with 200ng/mL EGF for 15min at 37°C in a media still containing the above inhibitors, then washed twice with 200μL per well of warmed DPBS and fixed in 150μL per well of 4% formaldehyde for 20min, then washed twice with 200μL per well 1x PBS then stored at 4°C in PBS containing 0.1U/μL RiboLock RNase Inhibitor until use on AVITI24 or prepared for bulk RNAseq by washing twice with 200uL per well warmed DPBS then lysed in 100μL per well TRI Reagent® and stored at −80°C until prepared for sequencing. On AVITI24, flowcells were sequenced using the Teton Human MAPK & Apoptosis Panel Kit (Element Biosciences, 830-00041) according to user guidance protocols.

### AVITI24 Teton™ Cytoprofiling

**AVITI24 Teton™ Cytoprofilng**
| **Components** | **Item Code** |
| --- | --- |
| Element AVITI24 System | 880-00004 |
| Teton Optimization Kit | 860-00022 |
| Teton Human MAPK & Cell Cycle Kit, 12 Well | 860-00023 |
| Teton Human MAPK & Apoptosis Kit, 12 Well | 860-00025 |
| Teton, Flow Cell Assembly Kit, 12 Well (2-pack) | 860-00028 |
| Teton Slide Kit, PLL - 12 Well (2-pack) | 860-00031 |
| Teton Slide Kit, Uncoated - 12 Well (2-pack) | 860-00032 |
| Teton Flow Cell Assembly Tool Set | 860-00033 |
| Teton Custom Add-On Protein Panel Assembly Kit | 860-00036 |

