## Supplementary Figures and Tables for "Unraveling Tyrosine-Kinase Inhibitor Resistance in NSCLC Cells via Same-cell Measurement of RNA, Protein, and Morphological Responses"

### Supporting Information

**Supplemental Table 1.** *Table of custom antibodies used in 55-plex protein panel, targeting MAPK/ERK pathway during TKI inhibition of NSCLC cells.*

| Target | Vendor | Catalog # |
|---|---|---|
| Cyclin D1 | Abcam | ab134175 |
| Phospho-Cyclin D1 | Cell Signaling Technology | 3300 |
| EGFR | Abcam | ab52894 |
| phospho-Y1068 EGFR | Abcam | ab40815 |
| SOS1 | Abcam | ab140621 |
| Ras | Abcam | ab52939 |

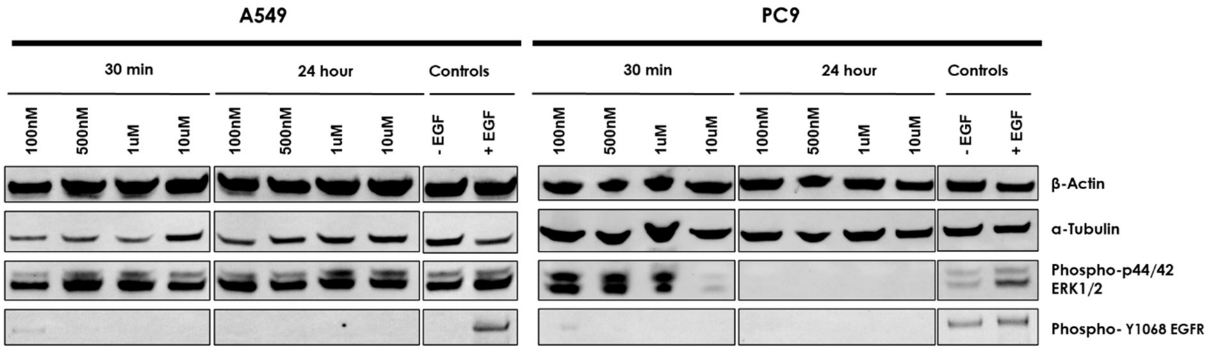


**Supplemental Figure 1.** *Western Blots of phospho-p44/42 ERK1/2 and phospho-Y1068 EGFR. A549 and PC9 cell lines treated with gefitinib at different doses either for 30 min or 24 hours demonstrating TKI gefitinib regulation of phospho-Y1068 EGFR at 10uM concentration of the drug in both cell lines. A549 was less responsive than PC9 which also demonstrated phospho-p44/42 ERK1/2 inhibition at 10uM concentration after 30-minute incubation and at all concentrations at 24 hours. Shown are targets for β-Actin and α-Tubulin for targets of phospho-p44/42 ERK1/2 and phospho-Y1068 EGFR.*

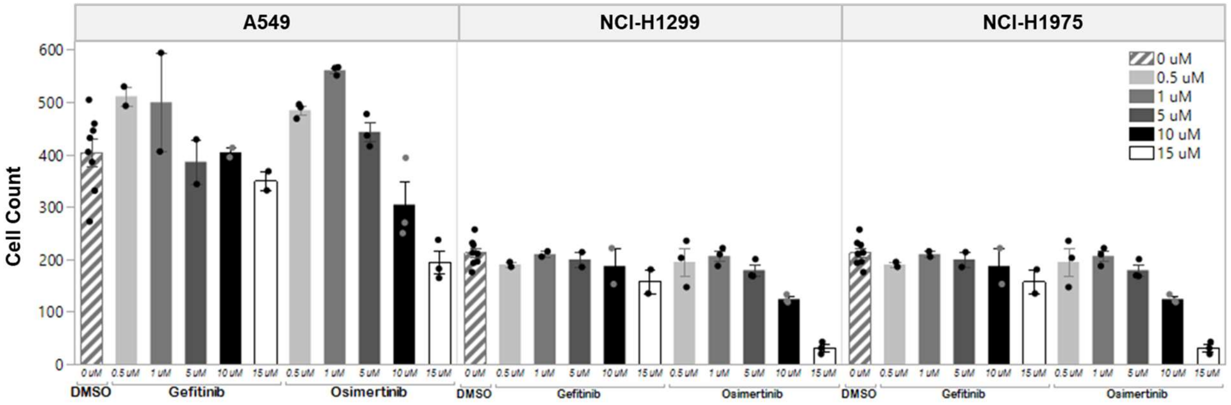


**Supplemental Figure 2.** *Cell count of A549, NCI-H1299, and NCI-H1975 after 24 hours of incubation with concentrations ranging from 0 to 15 uM of either gefitinib or osimertinib to determine sufficient cell response and cell retention for multiomic assay. 10uM concentration was determined to be the optimal condition that shows a reduced cell count compared to control in most cell lines while sustaining sufficient cells for multiomics on AVITI24 runs. Data was collected in triplicate and then averaged. Error bars indicate standard error.*

**A**

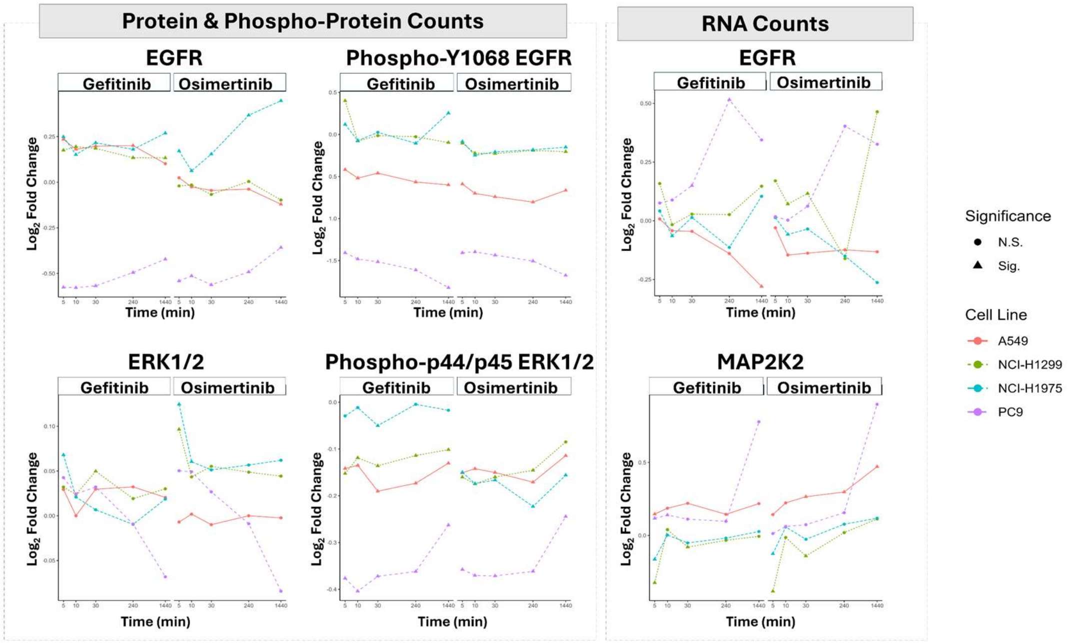


**B**

| Type | Target | Time (min) | A549 Gefitinb | A549 Osimertinib | NCI-H1299 Gefitinb | NCI-H1299 Osimertinib | NCI-H1975 Gefitinb | NCI-H1975 Osimertinib | PC9 Gefitinb | PC9 Osimertinib |
|---|---|---|---|---|---|---|---|---|---|---|
| Protein/Phospho Protein | EGFR | 5 | 0.235 | 0.024 | 0.175 | -0.021 | 0.249 | 0.171 | -0.576 | -0.541 |
| | | 10 | 0.181 | -0.026 | 0.195 | -0.015 | 0.151 | 0.062 | -0.578 | -0.513 |
| | | 30 | 0.198 | -0.045 | 0.186 | -0.067 | 0.216 | 0.154 | -0.568 | -0.562 |
| | | 240 | 0.201 | -0.038 | 0.134 | 0.004 | 0.180 | 0.367 | -0.495 | -0.492 |
| | | 1440 | 0.102 | -0.121 | 0.133 | -0.098 | 0.269 | 0.446 | -0.422 | -0.358 |
| | ERK1/2 | 5 | 0.029 | -0.007 | 0.032 | 0.097 | 0.068 | 0.125 | 0.043 | 0.050 |
| | | 10 | 0.000 | 0.002 | 0.024 | 0.043 | 0.021 | 0.060 | 0.024 | 0.049 |
| | | 30 | 0.030 | -0.010 | 0.050 | 0.055 | 0.007 | 0.051 | 0.032 | 0.027 |
| | | 240 | 0.032 | 0.000 | 0.019 | 0.049 | -0.010 | 0.057 | -0.009 | -0.009 |
| | | 1440 | 0.021 | -0.002 | 0.030 | 0.044 | 0.019 | 0.062 | -0.068 | -0.084 |
| | Phos-p44/p45 ERK1/2 | 5 | -0.142 | -0.151 | -0.152 | -0.160 | -0.029 | -0.150 | -0.376 | -0.358 |
| | | 10 | -0.135 | -0.142 | -0.119 | -0.175 | -0.012 | -0.174 | -0.404 | -0.370 |
| | | 30 | -0.190 | -0.150 | -0.136 | -0.160 | -0.050 | -0.167 | -0.372 | -0.371 |
| | | 240 | -0.173 | -0.171 | -0.114 | -0.146 | -0.005 | -0.223 | -0.362 | -0.362 |
| | | 1440 | -0.130 | -0.114 | -0.101 | -0.085 | -0.017 | -0.156 | -0.263 | -0.244 |
| | PhosY1068 EGFR | 5 | -0.418 | -0.589 | 0.400 | -0.104 | 0.119 | -0.084 | -1.407 | -1.407 |
| | | 10 | -0.519 | -0.700 | -0.078 | -0.223 | -0.074 | -0.248 | -1.481 | -1.396 |
| | | 30 | -0.460 | -0.741 | -0.014 | -0.227 | 0.025 | -0.207 | -1.514 | -1.435 |
| | | 240 | -0.565 | -0.804 | -0.027 | -0.190 | -0.105 | -0.182 | -1.611 | -1.505 |
| | | 1440 | -0.600 | -0.664 | -0.097 | -0.205 | 0.254 | -0.151 | -1.821 | -1.675 |
| RNA | EGFR | 5 | 0.008 | -0.030 | 0.158 | 0.170 | 0.042 | 0.013 | 0.075 | 0.017 |
| | | 10 | -0.042 | -0.146 | -0.017 | 0.071 | -0.065 | -0.058 | 0.088 | 0.002 |
| | | 30 | -0.045 | -0.138 | 0.028 | 0.116 | 0.014 | -0.035 | 0.150 | 0.062 |
| | | 240 | -0.140 | -0.124 | 0.026 | -0.161 | -0.114 | -0.150 | 0.516 | 0.403 |
| | | 1440 | -0.280 | -0.132 | 0.147 | 0.464 | 0.104 | -0.263 | 0.344 | 0.326 |
| | MAP2K2 | 5 | 0.147 | 0.143 | -0.324 | -0.382 | -0.163 | -0.126 | 0.117 | 0.013 |
| | | 10 | 0.186 | 0.223 | 0.040 | -0.014 | 0.003 | 0.058 | 0.140 | 0.061 |
| | | 30 | 0.220 | 0.265 | -0.080 | -0.141 | -0.051 | -0.027 | 0.111 | 0.074 |
| | | 240 | 0.145 | 0.297 | -0.033 | 0.019 | -0.019 | 0.077 | 0.097 | 0.156 |
| | | 1440 | 0.218 | 0.471 | -0.006 | 0.113 | 0.027 | 0.118 | 0.778 | 0.898 |

***Supplemental Figure 3.*** Protein, phospho-protein and RNA $\log_2$ fold change for indicated targets upon TKI treatment for four types of NSCLC cell lines A549, NCI-H1299, and NCI-H1975, and PC9 which indicate detection of clear regulation of key targets via AVITI24. **(A)** Plotted time course of $\log_2$ fold change versus time of TKI incubation. **(B)** Table summary of $\log_2$ fold change.

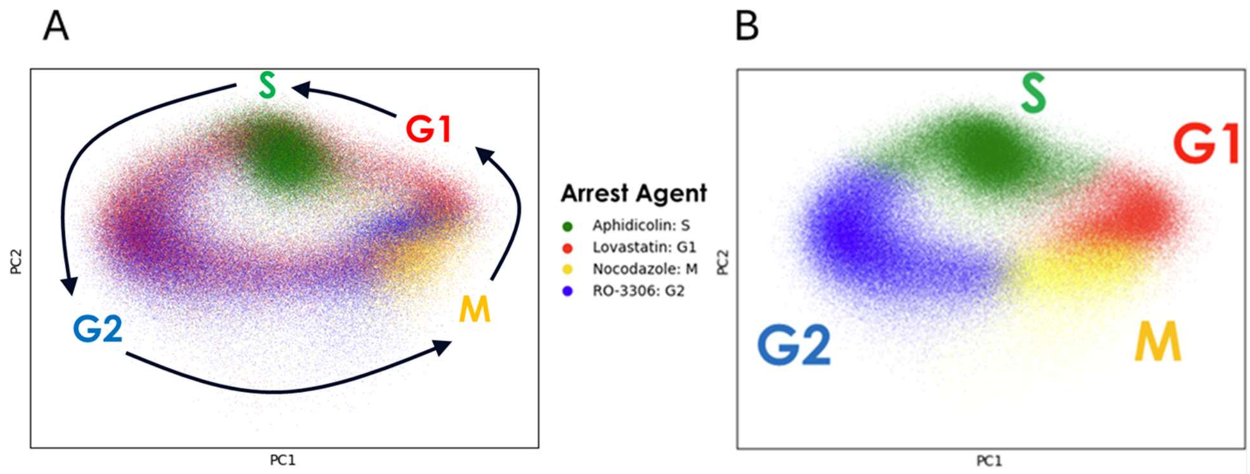


**Supplemental Figure 4.** *Cell cycle classifier training. A) RNA features projected to the top two principal components from two AVITI24 runs with different cell arrest agents in different wells showing enrichment of arrested cells in different regions of the observed anulus. Each dot represents a single-cell colored by the arrest agent applied to that well. B) Unsupervised clustering and visual association of each cluster to the enriched cell state based on all cells in A.*

**Supplemental Table 2.** *Quantified $log_2$ fold change values of CDCA3 and PLK1 at distinct cell phases during time course of gefitinib over 24 hours on A549, NCI-H1299, and NCI-H1975 cells.*

| | | CDCA3 | | | PLK1 | | |
|---|---|---|---|---|---|---|---|
| **Time** | **State** | **A549** | **NCI-H1299** | **NCI-H1975** | **A549** | **NCI-H1299** | **NCI-H1975** |
| **5** | **G1** | -0.0955 | 0.0242 | -0.0636 | -0.0541 | -0.0604 | -0.103 |
| | **G2** | -0.0932 | 0.0680 | 0.0468 | 0.0287 | -0.0398 | -0.0406 |
| | **M** | -0.0395 | 0.0405 | -0.0157 | -0.00274 | -0.0770 | -0.0994 |
| | **S** | -0.0661 | 0.0466 | -0.0193 | -0.0135 | -0.0354 | -0.117 |
| **10** | **G1** | -0.0303 | -0.00276 | -0.0254 | -0.000235 | 0.0264 | 0.00383 |
| | **G2** | -0.0159 | -0.00384 | -0.0578 | 0.0601 | 0.00812 | -0.0101 |
| | **M** | 0.0107 | -0.0107 | -0.0310 | 0.0443 | 0.0159 | -0.0246 |
| | **S** | -0.00899 | -0.0265 | -0.00790 | 0.0730 | -0.0334 | -0.00574 |
| **30** | **G1** | -0.0252 | -0.00987 | 0.0116 | -0.0199 | -0.00924 | 0.0182 |
| | **G2** | -0.0255 | -0.00865 | -0.00657 | 0.0264 | -0.0411 | -0.0212 |
| | **M** | -0.00114 | 0.00724 | -0.0460 | 0.0218 | -0.00811 | 0.0199 |
| | **S** | -0.0703 | -0.0184 | 0.00792 | -0.00254 | -0.0440 | -0.0316 |
| **240** | **G1** | 0.0411 | -0.162 | -0.0776 | -0.117 | -0.177 | -0.239 |
| | **G2** | -0.0183 | -0.273 | 0.0132 | -0.210 | -0.319 | -0.188 |
| | **M** | 0.0368 | -0.151 | -0.0911 | -0.0821 | -0.135 | -0.378 |
| | **S** | 0.0524 | -0.191 | 0.00562 | -0.173 | -0.269 | -0.355 |
| **1440** | **G1** | -0.146 | -0.132 | -0.339 | -0.286 | -0.119 | -0.553 |
| | **G2** | -0.465 | -0.760 | -0.480 | -0.463 | -0.762 | -0.916 |
| | **M** | -0.195 | -0.390 | -0.670 | -0.334 | -0.322 | -1.15 |
| | **S** | -0.188 | -0.230 | -0.469 | -0.281 | -0.486 | -0.912 |

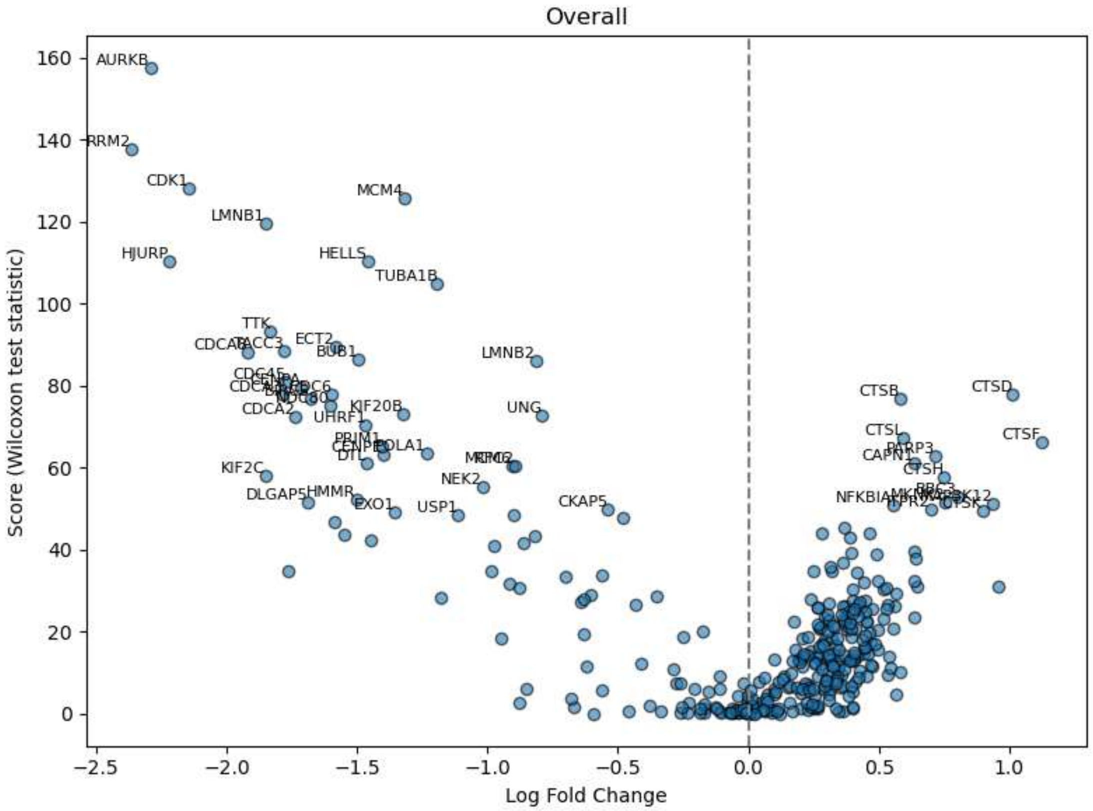


**Supplemental Figure 5.** *Differential expression statistics for Robustly regulated RNA targets in the A-state across 12 different AVITI24 runs. The minimum test statistic per target is shown. The top 20 targets that were robustly down-regulated are: NDC80, BIRC5, CDCA3, CDC6, CENPA, CDC45, LMNB2, BUB1, CDCA8, TACC3, ECT2, TTK, TUBA1B, HELLS, HJURP, LMNB1, MCM4, CDK1, RRM2, and AURKB. The top 20 targets that were robustly up-regulated are: CTSD, CTSB, CTSL, CTSF, PARP3, CAPN1, CTSH, BBC3, MKNK2, MAP3K12, NFKBIA, ITPR2, CTSK, XIAP, BAK1, CTSZ, PARP4, CFLAR, CASP7, and TRADD.*

A

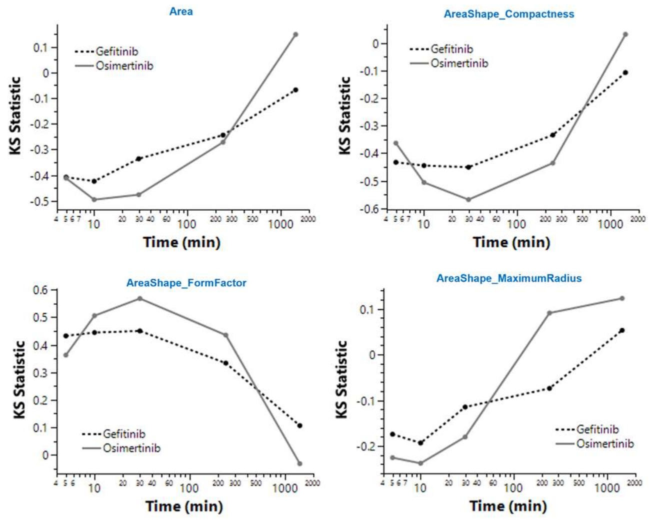
Area
AreaShape_Compactness
AreaShape_FormFactor
AreaShape_MaximumRadius
Gefitinib
Osimertinib
KS Statistic
Time (min)

B

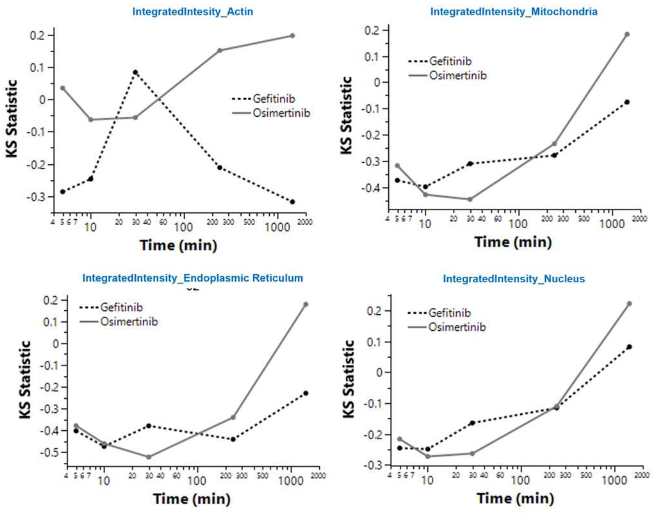
IntegratedIntesity_Actin
IntegratedIntensity_Mitochondria
IntegratedIntensity_Endoplasmic Reticulum
IntegratedIntensity_Nucleus
Gefitinib
Osimertinib
KS Statistic
Time (min)

C

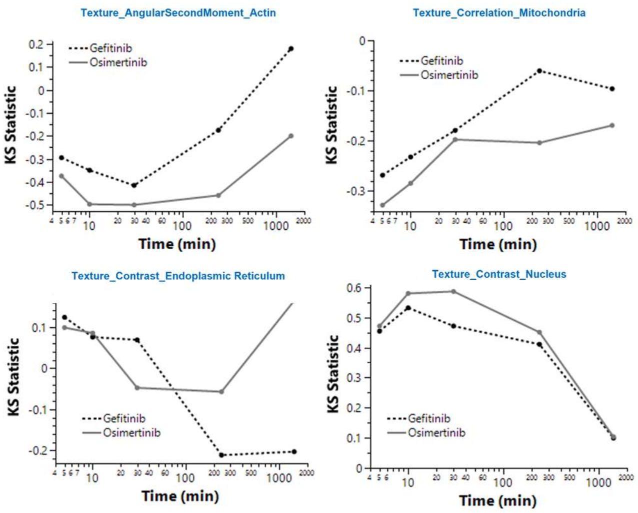
Texture_AngularSecondMoment_Actin
Texture_Correlation_Mitochondria
Texture_Contrast_Endoplasmic Reticulum
Texture_Contrast_Nucleus
Gefitinib
Osimertinib
KS Statistic
Time (min)

**Supplemental Figure 6.** *Affected cell paint morphology features during 24-hour time course of 10μM Gefitinib and Osimertinib treatment on PC9 cells. Plotted are Kolmogorov-Smirnov (KS) statistic values, representing the amount of deviation from the untreated EGF activated control distribution versus time of TKI incubation. Representative area-related metrics (A), cell paint intensity (B), and texture-related metrics (C) are shown.*
